# Field manipulation of competition among hybrids reveals dynamic and highly stable features of a complex fitness landscape driving adaptive radiation

**DOI:** 10.1101/756908

**Authors:** Christopher H. Martin, Katelyn Gould

## Abstract

The effect of the environment on fitness in natural populations is a fundamental question in evolutionary biology. However, experimental manipulations of environment and phenotype are rare. Thus, the relative importance of the competitive environment versus intrinsic organismal performance in shaping the location, height, and fluidity of fitness peaks and valleys remains largely unknown. We experimentally tested the effect of competitive environment on the fitness landscape driving the evolution of novelty in a sympatric adaptive radiation of a generalist and two trophic specialist pupfishes, a scale-eater and molluscivore, endemic to San Salvador Island, Bahamas. We manipulated phenotypes, by generating 2,611 F4/F5 lab-reared hybrids, and competitive environment, by altering frequencies of rare phenotypes between high- and low-frequency field enclosures, then tracked hybrid survival in two natural lake populations on San Salvador. We found no evidence of frequency-dependent effects on survival fitness landscapes, indicating robustness to the competitive environment. Although survival surfaces favored alternate phenotypes between lakes, joint fitness estimation across lake environments supported multiple fitness peaks for generalist and molluscivore phenotypes and a large fitness valley isolating the most divergent scale-eater phenotype, strikingly similar to a previous independent field experiment. The consistency of this complex fitness landscape across competitive environments, multivariate trait axes, and spatiotemporal heterogeneity provides surprising evidence of stasis in major features of fitness landscapes despite substantial environmental variance, possibly due to absolute biomechanical constraints on diverse prey capture strategies within this radiation. These results challenge competitive speciation theory and highlight the interplay between organism and environment underlying static and dynamic features of the adaptive landscape.

## Introduction

The adaptive landscape, the complex mapping of fitness onto phenotype or genotype, is both a central unifying concept in evolutionary biology and an empirical measurement (Wright 1932; Lande 1979; Gavrilets 2004; Carneiro & Hartl 2010; Svensson & Calsbeek 2012) which links the microevolutionary processes of natural and sexual selection in wild populations with macroevolutionary patterns of speciation, novelty, and adaptive radiation (Lande & Arnold 1983; Arnold *et al.* 2001; Kingsolver *et al.* 2001; Martin & Richards 2019).

Despite its central importance, it remains unclear what factors shape the fitness landscape across space and time. In classical views arising from both Wright’s (Wright 1932) and Simpson’s (Simpson 1944a) original conceptions of genotypic and phenotypic fitness landscapes, respectively, and Fisher’s geometric model (Fisher 1930), fitness optima occur on a high-dimensional landscape with both static and dynamic features. Stability of fitness optima is due to negative epistasis within genotypic networks (Whitlock *et al.* 1995; Weinreich *et al.* 2005) and functional tradeoffs between different ecological niches or collections of similar niches known as adaptive zones (Simpson 1944b; Higham *et al.* 2016). A more recent paradigm originating from game theory (Abrams *et al.* 1993; Abrams 2001; Bolnick 2004; Doebeli & Dieckmann 2005) emphasizes that the fitness landscape is dynamic and resembles a trampoline: as the relative frequency of phenotypically similar individuals increases, their fitness decreases due to increased competition for resources, whereas rare phenotypes at the extremes of the normal distribution of phenotypes within a population have a fitness advantage (Gavrilets 2004). This is known as negative frequency-dependent disruptive selection and can lead to ecological speciation in sympatry, even while adapting to a unimodal resource distribution (Dieckmann & Doebeli 1999; Matessi *et al.* 2002; Doebeli *et al.* 2005; Bürger & Schneider 2006; Otto *et al.* 2008). Laboratory and field studies of natural populations provide extensive support for negative frequency-dependent disruptive selection (Pfennig 1992; Hori 1993; Sinervo *et al.* 2000; Schluter 2003; Bolnick 2004; Kassen *et al.* 2004; Olendorf *et al.* 2006; Bolnick & Lau 2008; Weeks & Hoffmann 2008; Koskella & Lively 2009; Svensson & Calsbeek 2012; Kusche *et al.* 2012; Bolnick & Stutz 2017; Nosil *et al.* 2018), to the extent that some investigators assert its universality in all natural populations (Haller & Hendry 2014). Due to its elegance and mathematical tractability, frequency-dependence has also been widely adopted by theorists as the sole mechanism for disruptive selection in most speciation models (Gavrilets 2004; Polechová & Barton 2005; Martin & Richards 2019).

However, the relative contributions of the competitive environment (i.e. frequency-dependence) versus static fitness optima (i.e. functional tradeoffs) in shaping the broader topography of fitness landscapes across multiple species remains unclear, despite the importance of these interactions in providing the bridge between microevolutionary processes within a population and macroevolutionary scale patterns of diversification and adaptive radiation (Arnold *et al.* 2001; Martin & Richards 2019). For example, although negative frequency-dependent disruptive selection may be ubiquitous within populations, the phenotypic similarity necessary for individuals to compete with each other within the same ecological niche is rarely measured (i.e. the competition kernel), and can have major impacts on speciation (Dieckmann & Doebeli 1999; Baptestini *et al.* 2009). Similarly, shifts in the location of a fitness optimum due to environmental stochasticity are often assumed to follow a Brownian motion process (Grant & Grant 2002; Hansen *et al.* 2008) without accounting for hard boundaries imposed by functional tradeoffs or biophysical constraints (but see: (Enquist & Niklas 2002; Boucher & Demery 2016)). Conversely, at broader macroevolutionary timescales the role of stable fitness optima in shaping trait diversification across a radiation is frequently tested and supported by fitting Ornstein-Uhlenbeck models of trait diversification using phylogenetic comparative methods (Butler & King 2004; Harmon *et al.* 2010; Beaulieu & O’Meara 2012; O’Meara 2012; Uyeda & Harmon 2014) while not accounting for the effects of density- and frequency-dependent selection due to competition within a community (but see (Rabosky 2009; Harmon *et al.* 2019; Landis *et al.* 2019)).

Previous field experimental tests of frequency-dependent selection usually focused on only a single species or pair of ecomorphs (Pfennig 1992; Sinervo *et al.* 2000; Schluter *et al.* 2003; Bolnick 2004; Olendorf *et al.* 2006; Weeks & Hoffmann 2008; Calsbeek *et al.* 2009; Kusche *et al.* 2012; Bolnick & Stutz 2017) and there are still few studies spanning multiple habitats, traits, time periods, or species. In the few studies at these larger scales, rare transgressive hybrid phenotypes appear to suffer a fitness cost, not an advantage (Keagy *et al.* 2015; Martin 2016a). For example, in a hybrid mesocosm experiment investigating male-male competition in stickleback, the rarest transgressive phenotypes experienced the lowest reproductive success (Keagy *et al.* 2015), in contrast to predictions of sexual selection as a diversifying force ((Seehausen & Schluter 2004); but see (Servedio & Burger 2014; Kopp *et al.* 2017)). We previously estimated multiple fitness peaks driving adaptive radiation in the San Salvador Island pupfishes by measuring the growth and survival of laboratory-reared F2 intercross and backcross hybrids placed in high- and low-density field enclosures (Martin & Wainwright 2013a). Hybrid phenotypes resembling the widespread generalist species were isolated by a local fitness peak, separated by a fitness valley from a higher fitness peak corresponding to hybrid phenotypes resembling the molluscivore specialist, whereas hybrid phenotypes resembling the scale-eating specialist suffered the lowest growth and survival. Interestingly, rare phenotypes in this experiment did not experience a survival advantage and the scale of competitor frequency-dependent survival appeared to operate only within the range of hybrid phenotypic diversity observed in the generalist population, not across fitness valleys among all three species (Martin 2016a). These few experimental studies spanning multiple species suggest that negative frequency-dependent selection may primarily operate within a population, rather than between species occupying highly divergent generalist and specialist ecological niches; however, experimental tests manipulating the frequency of rare hybrid phenotypes are needed to investigate the static and dynamic features of fitness landscapes.

Here we experimentally manipulated the frequency of rare hybrid phenotypes to test for an effect on the stability of fitness optima in a tractable system for empirical measurements of the fitness landscape during nascent adaptive radiation. We generated transgressive hybrid phenotypes by multiple rounds of backcrossing and intercrossing between a generalist and two trophic specialist *Cyprinodon* pupfish species in two independent hybrid populations, manipulated the frequency of rare hybrid phenotypes between treatments in two independent lake populations on San Salvador Island, Bahamas, and then tracked individual hybrid survival and growth rates. This experimental procedure creates a normally distributed but highly variable distribution of hybrid phenotypes recreating the early stages of diversification before adaptive radiation began on San Salvador Island. We found negligible effects of competitor frequency on the survival of hybrids in both lakes despite major differences in the topography of fitness landscapes. However, all four fitness surfaces supported multiple fitness peaks for generalists and molluscivore phenotypes combined with a large region of low fitness isolating the novel scale-eating specialist, highly similar to survival patterns in a previous independent field experiment. These results provide strong empirical field support for both environment-dependent fitness dynamics across space and time and environment-independent stasis of major feature of a complex fitness landscape, which help explain the rare evolution of the scale-eating ecological niche specialist.

## Methods

### Frequency-manipulation within field enclosures

We individually tagged and photographed 2,611 F4-F5 outbred juvenile hybrids resulting from crosses of all three species, generalist (*C. variegatus*), molluscivore (*C. brontotheroides*), and scale-eater (*C. desquamator*), from two different isolated lake populations on San Salvador Island, Bahamas before release into a high- and low-frequency field enclosure in each lake (Fig. 1; Table S1). Frequency-manipulation increased the frequency of transgressive hybrid phenotypes in the high-frequency treatment and generalist-type hybrids in the low-frequency treatment, resulting in a significant reduction in phenotypic variance on discriminant axis 2 (predominantly nasal protrusion) in lake 1 (Levene’s test, *P* < 0.0001) and discriminant axis 1 (predominantly oral jaw size) in lake 2 (Levene’s test, *P* < 0.0001) within the bivariate discriminant morphospace separating all three parental species. The total density of hybrids was held approximately constant between high and low-frequency treatments (lake 1: high/low: 923/823 individuals; lake 2: high/low: 842/819 individuals; Table S1).

**Fig. 1.**
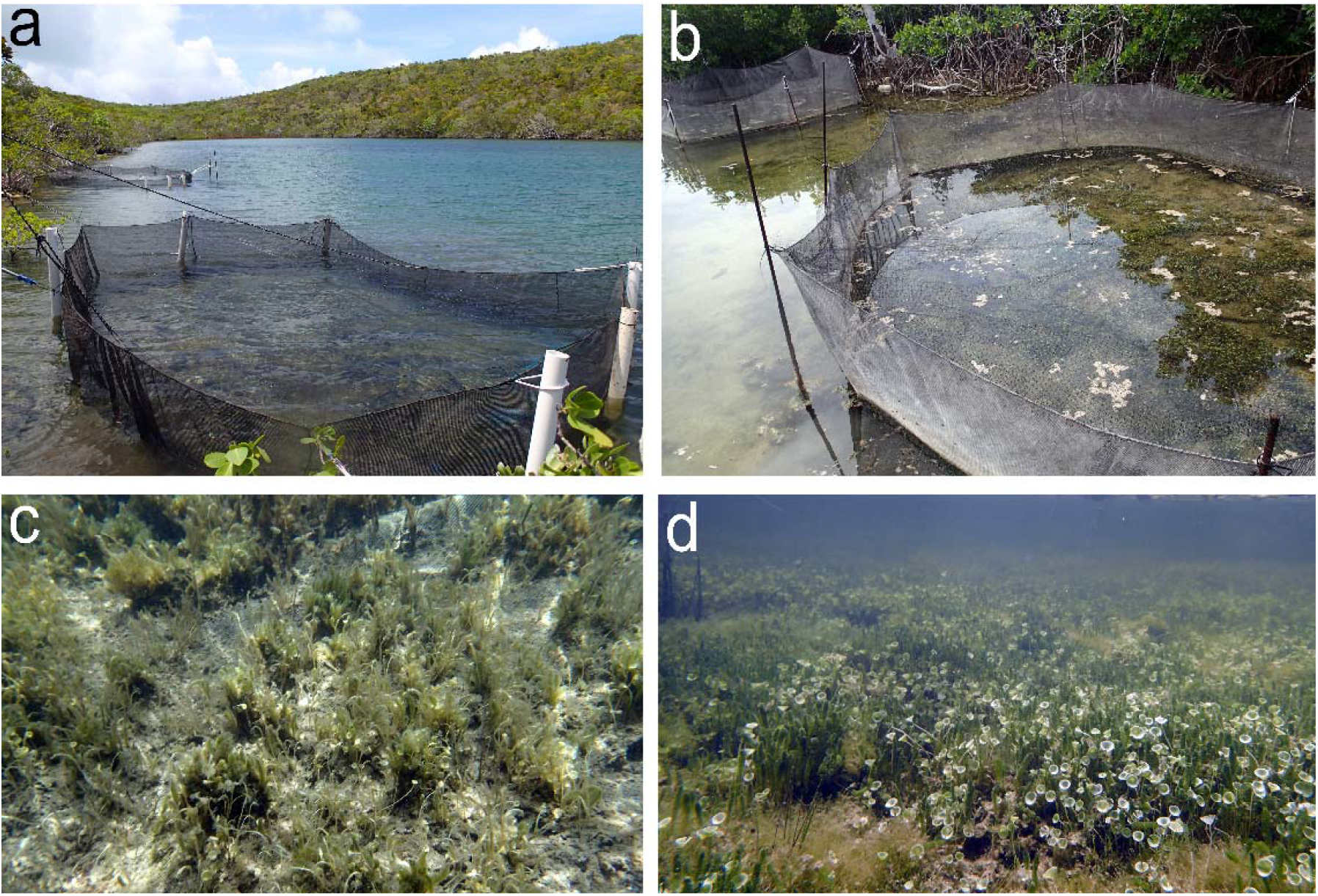
**High- and low-frequency field enclosures** and the associated benthic macroalgae communities inside each enclosure typical of surrounding littoral zone habitats in lake 1 (*a,c*: Crescent Pond) and lake 2 (*b,d*: Little Lake) after 3 month and 11 month field exposure periods, respectively.

### Survival in field enclosures and laboratory control environments

To sample from a broader range of environmental variability, we measured hybrid survival after 3 months in lake 1 (high-frequency: 77.1% survival; low-frequency: 75% survival) and after 11 months in lake 2 (high-frequency: 1.4% survival; low-frequency: 1.2% survival; Table S1), in the latter case spanning most of the adult lifespan (Martin *et al.* 2016) but avoiding mortality due to senescence. There were no differences in survival probability between treatments in each lake (two-way logistic regression, treatment effect: *P* = 0.237). For a control comparison, additional hybrids from each lake population (*N* = 199 total individuals) were simultaneously tagged, raised in laboratory aquaria, and their deaths and growth rates were tracked over one year concurrently with field experiments.

### Morphometrics

Phenotypic similarity of each hybrid to the three parental species in each lake (*n* = 236) was calculated from 30 linear traits and angles measured from three pre-release photographs of each fish (Fig. S2). These traits were used to estimate two linear discriminant (LD) axes with major loadings (Table S2) of oral jaw size (LD axis 1) and nasal protrusion (LD axis 2), diagnostic traits of each specialist species and major axes of rapid trait diversification within this radiation (Martin & Wainwright 2011; Martin 2016b). Indeed, after correcting for standard length, residual jaw length variation within our hybrid populations exceeded the range of variation observed across allopatric *Cyprinodon* species and outgroup Cyprinodontidae species spanning over 20 million years since their most recent common ancestor (data from (Martin & Wainwright 2011); Fig. 2). Further details on rearing conditions, field enclosures, morphometrics, and statistical analyses are provided in the supplemental methods.

**Fig. 2.**
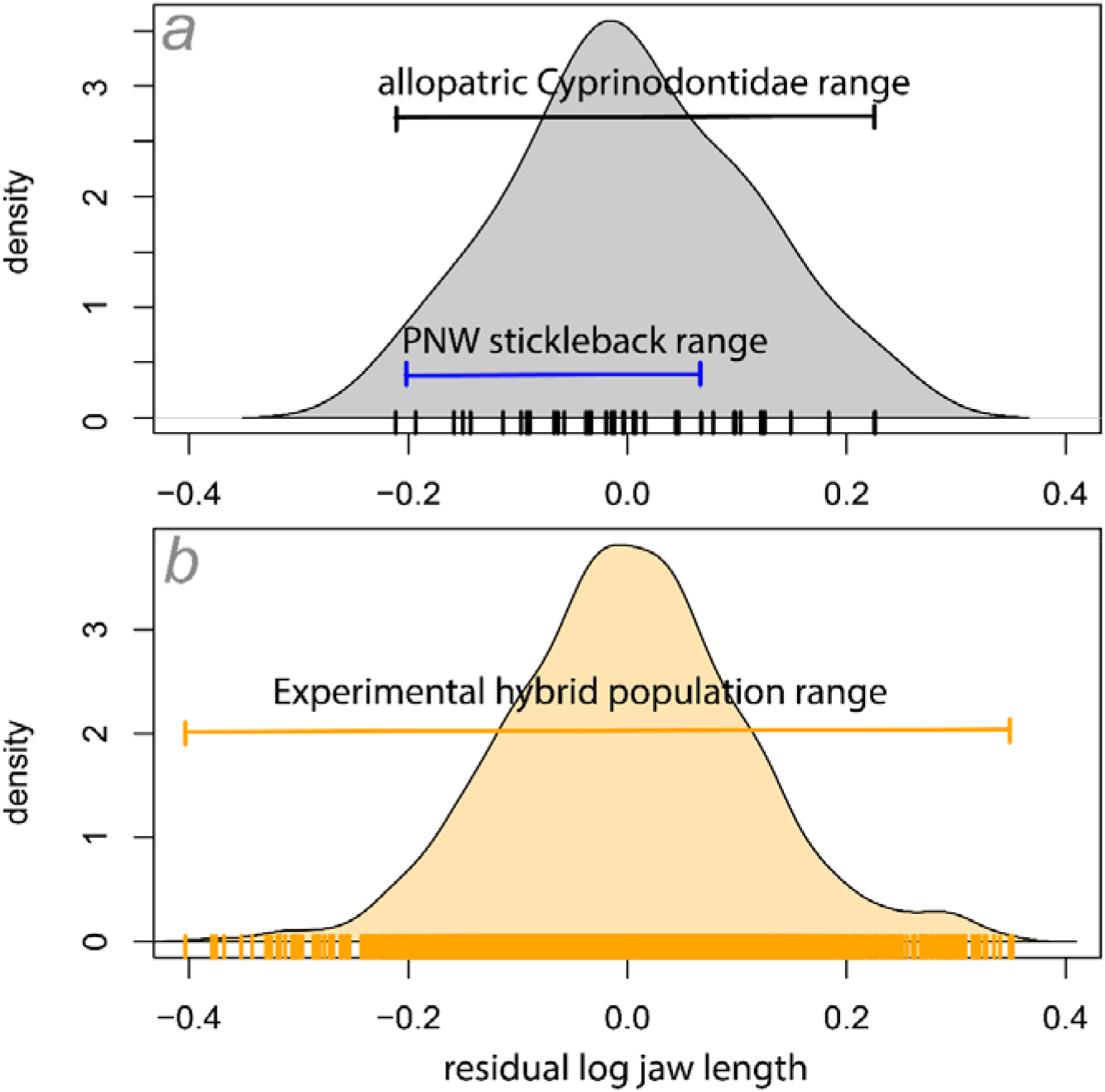
**Residual jaw length variation** in *a)* **allopatric Cyprinodontidae species** (grey) diverging over 20 million years ago ((Furness & Reznick 2015); data from (Martin & Wainwright 2011)) and *b)* lab-reared F4/F5 hybrid populations (orange) measured from the field experiment in this study. The minimum and maximum residual upper jaw lengths across allopatric stickleback populations in the Pacific Northwest (PNW; data from Fig. 1 reported in (Lavin & McPhail 1986)) are also included for comparison. Each *a)* species mean or *b)* individual F4/F5 hybrid is represented as tick marks on the x-axis plus density plots as an estimate of residual variation. Residuals were calculated from a linear regression of log-transformed lower jaw length on log-transformed standard length without mean and variance standardization for comparison on the same absolute scale across all three studies.

## Results

### Strong directional and nonlinear selection for generalist and molluscivore phenotypes

We fit thin-plate splines using generalized cross-validation or restricted maximum likelihood to survival data to visualize fitness landscapes across the discriminant morphospace (Nychka *et al.* 2017). We found evidence for directional selection on the survival of hybrid phenotypes most similar to generalist and molluscivore phenotypes in lake 1 (Fig. 3). Despite low survival rates after 11 months in lake 2, we found strong evidence of nonlinear divergent selection for hybrid phenotypes resembling the molluscivore (Fig. 3). These landscapes were estimated from very few survivors (*n* = 22); however, the density of non-survivors in the surrounding regions of morphospace provides robust support for nonlinear divergent selection for the molluscivore phenotype in this lake. Patterns of survival in the wild were contrasted by strong directional selection for hybrids resembling the scale-eater in laboratory control populations from both lakes (Fig. 3).

**Fig. 3.**
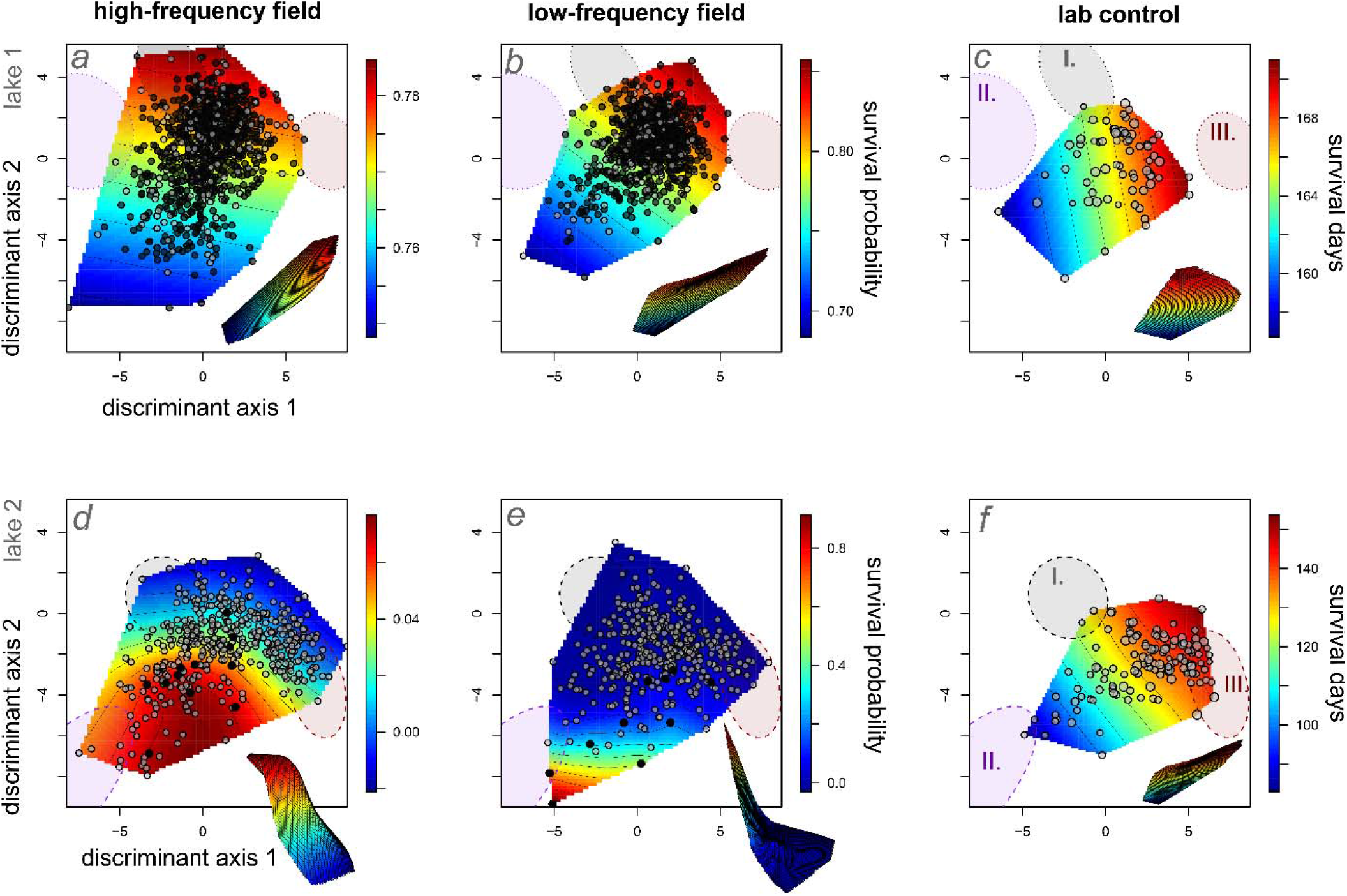
Survival fitness landscapes for hybrid populations in high- (first column) and low-frequency (second column) field enclosures and laboratory controls (third column). Thin-plate splines predict the probability of survival (heat color) across a single linear discriminant morphospace separating generalist and scale-eater phenotypes (x-axis: LD1) and generalist and molluscivore phenotypes (y-axis: LD2). Survivors in field enclosures are depicted in black relative to deaths over the 3-month and 1-year exposure periods, respectively. Laboratory control points are proportional to the number of days each hybrid survived within 151-liter aquaria. All hybrids are plotted within a shared linear discriminant morphospace for parental species’ phenotypes calculated from lab-reared F1 individuals of all three parental populations from both lakes (first row: lake 1; second row: lake 2). 95% confidence ellipses for each parental population in each lake are shown for generalists (I. grey), molluscivores (II. purple), and scale-eaters (III. red); note that molluscivores in lakes 1 and 2 show large differences in mean nasal protrusion distance (Martin & Feinstein 2014) leading to their separation along discriminant axis 2. Survival surfaces are depicted in three dimensions in the offset images.

We also used generalized projection pursuit regression to estimate the two multivariate linear phenotypic axes most strongly associated with survival across the 30-trait morphospace (Tables S3-S4) without making unfounded parametric assumptions about quadratic curvature as in canonical rotation analyses (Mitchell-Olds & Shaw 1987; Schluter & Nychka 1994; Morrissey 2014). Thus, projection pursuit regression estimates composite multivariate splines maximizing fitness surface curvature in the 30-trait space and is the nonlinear analog of the quadratic canonical rotation approach. Visualization of survival fitness landscapes on the first two major axes of nonlinear selection indicated that the most transgressive hybrid phenotypes (i.e. least similar to parental phenotypes) suffered the lowest survival probability across treatments in both lakes (Fig. 4). In contrast, survival in laboratory control populations was shifted or opposite to the direction of selection in field enclosures along these major axes of selection (Fig. 4).

**Fig. 4.**
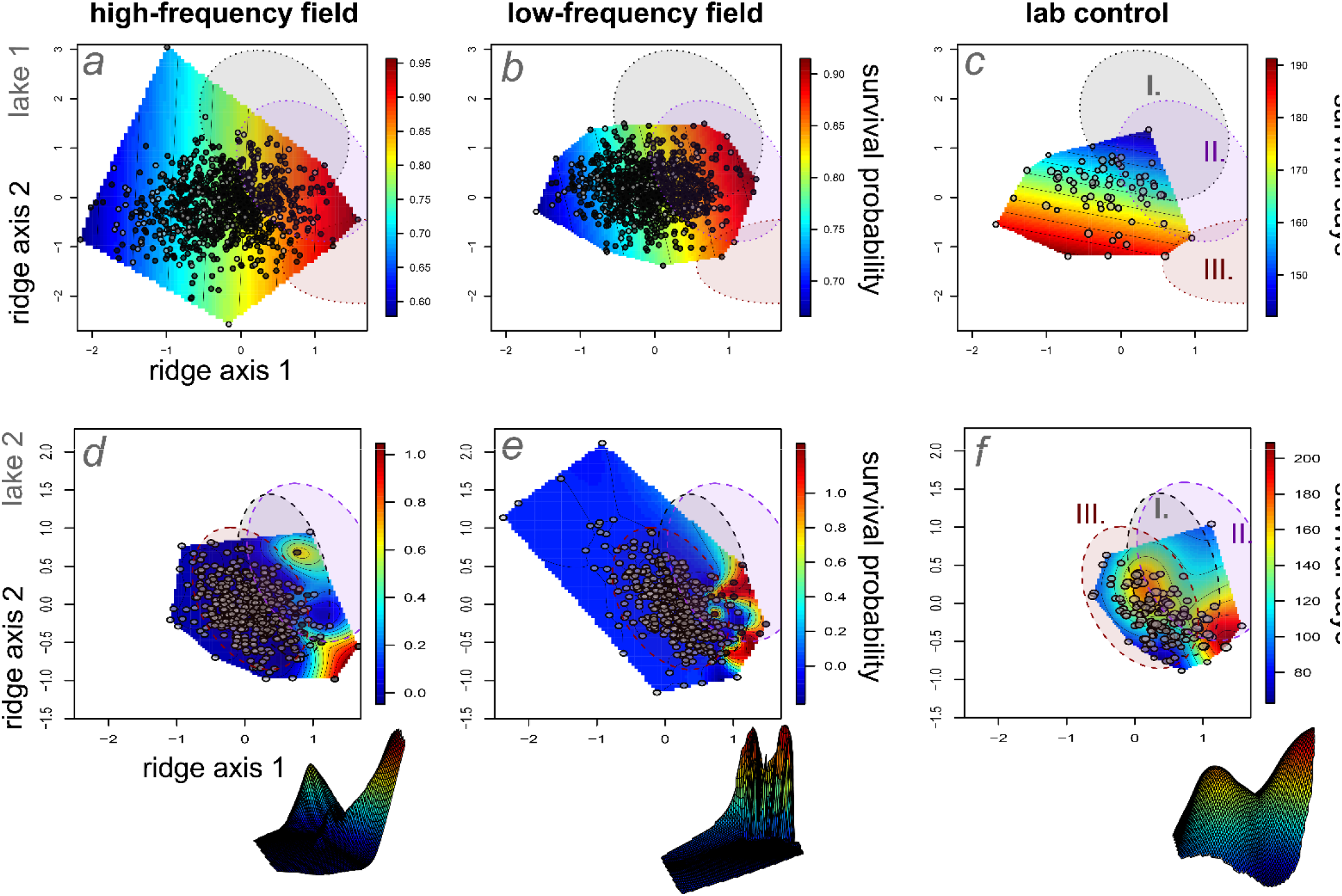
Survival fitness landscapes for hybrid populations in high- (first column) and low-frequency (second column) field enclosures and laboratory controls (third column) across the two major ridge axes of nonlinear selection within the 30-trait morphospace. Thin-plate splines predict the probability of survival (heat color) across the two major ridge axes associated with survival (Table S3-4), estimated separately for lake 1 (first row) and lake 2 (second row) hybrid populations using generalized projection pursuit regression (Tables S3-S4). Survivors in field enclosures are depicted in black relative to deaths over the 3-month (first row) and 1-year (second row) exposure periods, respectively (Table S1). Laboratory control points (third column) are proportional to the number of days each hybrid survived within 151-liter aquaria. 95% confidence ellipses for each parental population in each lake are shown for generalists (I. grey), molluscivores (II. purple), and scale-eaters (III. red). Nonlinear survival surfaces are depicted in three dimensions in the offset images.

We estimated the strength of multivariate selection gradients along these two major axes of selection and found significant evidence of directional selection (*P* < 0.00001) on ridge axis 1 in both lakes and marginal evidence of directional selection on ridge axis 2 in lake 2 (Table S5). The traits with the highest loadings on ridge axis 1 were a) lower jaw length and b) distance from the jaw joint to the orbit and on ridge axis 2 were a) angle between the premaxilla and orbit and b) distance from the premaxilla to the pectoral girdle (Tables S3-S4), further supporting strong selection on craniofacial trait diversification within this radiation (Martin 2016b).

### No evidence of frequency-dependent survival differences between treatments

We found no evidence of significant treatment effects on either survival probability or the overall topography of the survival fitness landscape within the discriminant morphospace (Table S6). The fixed effect of treatment did not improve the fit to the survival data in any of the generalized additive models examined. Instead, models without the effect of frequency treatment were strongly favored (Table S6; ΔAIC = 11). Similarly, we found no evidence for frequency-dependent effects on survival between treatments on the two major axes of nonlinear selection in either lake estimated from generalized projection pursuit regression (Table S7). Models without the effect of treatment provided a marginally better fit to the survival data on the two major axes of nonlinear selection in lake 1 (ΔAIC = 1) and were supported in lake 2 (Table S7; ΔAIC = 1.8).

### Phenotypic scale of frequency-dependence for growth rate but not survival

Within each enclosure we estimated the Mahalanobis distance for each hybrid individual, the distance to the mean hybrid phenotype in the 30-dimensional morphospace correcting for trait covariances, as an estimate of the rarity of each individual phenotype. We also calculated the Euclidean nearest neighbor distance to the ten most similar hybrid phenotypes in 30-dimensional morphospace for each hybrid as an estimate of the local frequency of competing phenotypes. Overall, these measures estimate the frequency of similar hybrid phenotypes relative to each hybrid’s phenotype to examine the phenotypic scale of frequency-dependence within each enclosure (Martin 2016a). The frequency of similar phenotypes was not significantly associated with residual variation in survival not explained by hybrid phenotype along the two discriminant axes (Fig. S4; Table S6). Generalized additive models including the fixed effect of competitor frequency (distance to mean phenotype) only marginally improved the fit to the survival data (Table S6; ΔAIC = 1; similar results were found when substituting nearest neighbor Euclidean distance for Mahalanobis distance: Fig. S5).

In contrast, generalized additive models including the fixed effect of competitor frequency (both Mahalanobis and nearest neighbor distance) strongly improved the fit to the growth data in lake 1 within the discriminant morphospace, even after accounting for the treatment effect (Table S6, Fig. S6; ΔAIC = 12; lake 2 was excluded from all growth rate analyses due to the low number of survivors). Similarly, for the two major axes of nonlinear selection estimated from generalized projection pursuit progression, models including the fixed effect of competitor frequency substantially improved the fit to the growth data, even after accounting for the treatment effect (Table S7; ΔAIC = 3.85).

### Joint inference of a single fitness landscape indicates strong similarity to a previous independent field experiment

Generalized additive modeling enables estimation and visualization of a joint fitness landscape after controlling for the effects of lake/exposure period and treatment. The best supported models for survival included a fixed effect of lake and no effect of treatment, with either two univariate smoothing splines or a thin-plate spline and two smoothing splines modeling selection within the discriminant morphospace (Table 2). Models including spline terms were strongly supported over models including fixed linear effects of the discriminant axes (Table 2; ΔAIC = 18.7).

Strikingly, the best combined model for survival (across all four treatments in both lakes) supported an isolated fitness peak for hybrids resembling the generalist separated by a fitness valley from a region of higher fitness corresponding to the molluscivore phenotype (Fig. 5), similar to a previous field experiment in 2011 using F2 hybrids in these same lakes (Martin & Wainwright 2013a). However, in the previous experiment few of the F2 hybrids fell within the phenotypic range of lab-reared scale-eaters (Martin & Wainwright 2013a). In our current experiment, over 70 hybrids occurred within the 95% confidence ellipse of lab-reared F1 scale-eater phenotypes within the discriminant morphospace. This provides evidence for environment-independent features of a complex fitness landscape across years, seasons, divergent lake environments, and experimentally-manipulated competitor frequencies.

**Fig. 5.**
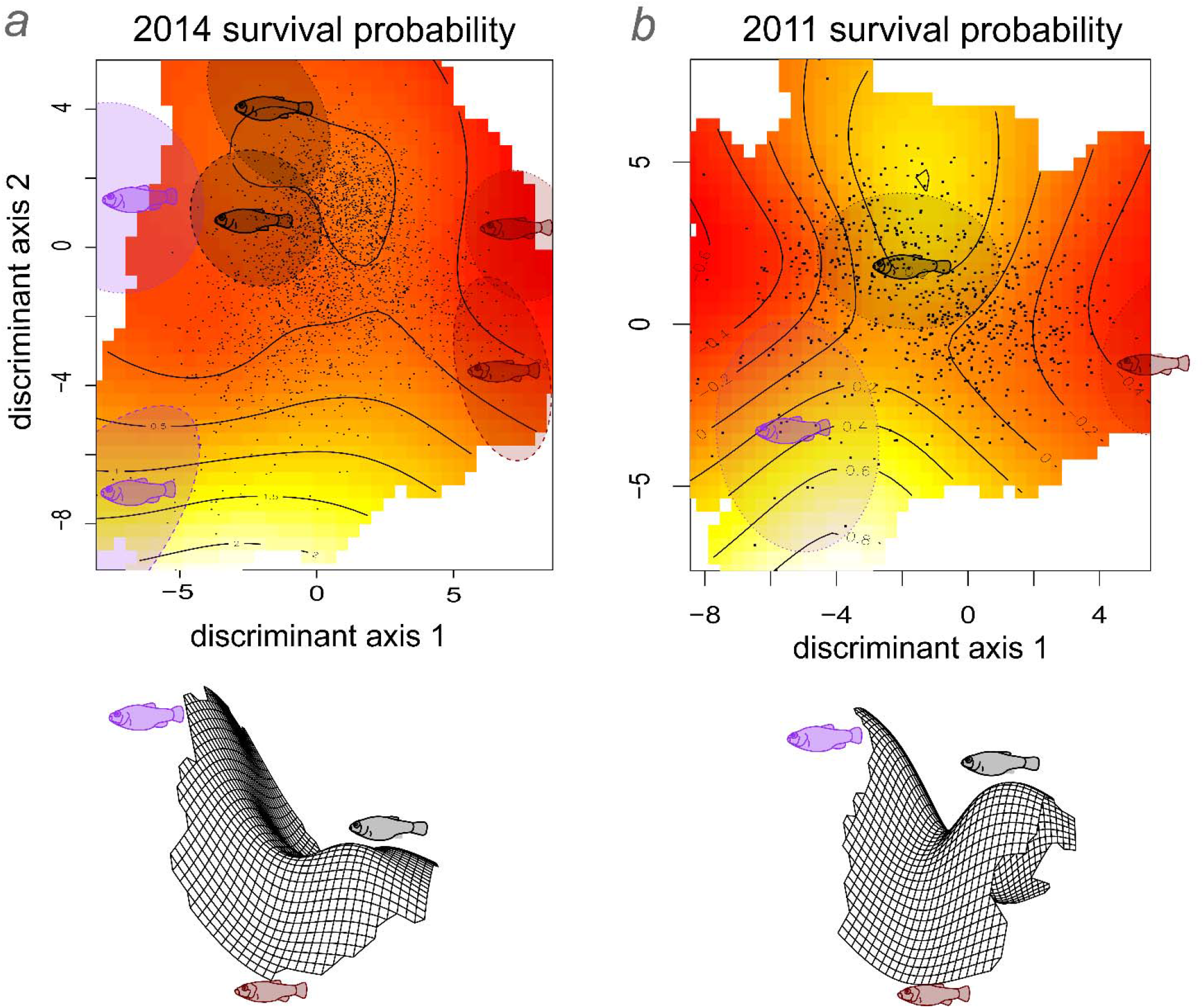
Survival fitness landscapes compared between independent field experiments in *a)* 2014/2015 (this study) and *b)* 2011 estimated from thin-plate splines using general additive models. Thin-plate splines in two- and three-dimensions estimate hybrid survival probability *a)* integrating across lake/field exposure period and high/low-frequency treatment for all four enclosures in the 2014-2015 field experiment based on the best-fitting generalized additive model (Table S6) and *b)* in only the high-density field enclosure in lake 1 from the 2011 field experiment (Martin & Wainwright 2013a), calculated from the original data using a generalized additive model. Thin-plate splines are depicted within the linear discriminant morphospace separating generalist and scale-eater phenotypes (x-axis: LD1) and generalist and molluscivore phenotypes (y-axis: LD2) calculated from lab-reared individuals of all three parental populations in both lakes (note different mean phenotypes for parental species between lake 1 and lake 2 in panel a). 95% confidence ellipses show the location of generalist (grey), molluscivore (purple), and scale-eater (red) parental populations from lake 1 (small dashed line) and lake 2 (large dashed line). Discriminant morphospaces were estimated from independent sets of phenotypic measurements in each panel. The 2014-2015 morphospace was estimated from 30 traits from six independent parental populations, three from each lake; the 2011 morphospace was estimated from a set of 16 traits (partially overlapping with the 30-trait set) from the three lab-reared parental populations from lake 1 (Crescent Pond).

It is possible that some regions of the high-dimensional trait space may still connect scale-eater phenotypes to other regions of the morphospace through a fitness ridge (Gavrilets 1997, 1999). To further explore the relative fitness of scale-eater hybrids, we visualized selection across all directions in the 30-trait morphospace by repeatedly sampling a random subset of 15 traits, calculating a discriminant axis for scale-eaters relative to generalists within this subspace, and estimating a survival spline for hybrid phenotypes on each arbitrary multivariate axis (Fig. 7). This results in a visualization of all possible fitness paths between generalist and scale-eater hybrid phenotypes for all subspaces within the 30-trait morphospace and, importantly, aligns these multivariate linear axes in the same direction from generalist to scale-eater phenotype in order to compare survival curves across random subsets of the trait data. In three out of four field enclosures (with no relationship in the fourth), hybrids resembling scale-eaters suffered the lowest survival across nearly all visualized fitness paths, supporting their position in a high-dimensional fitness valley within our morphospace (Fig. 7). These analyses suggest a robust fitness minimum within the adaptive landscape isolating the scale-eater phenotype from other species.

## Discussion

### No evidence of frequency-dependent survival in a multi-peak fitness landscape

We conducted an experimental field test of frequency-dependent selection in a nascent adaptive radiation of trophic specialist pupfishes. We found negligible evidence of frequency-dependent survival between treatments manipulating the frequency of rare transgressive hybrid phenotypes in two independent lake environments and hybrid populations nor any relationship between survival and the frequency of competitors (Figs. 3-5, S4-S5; Tables S6-S7). These patterns were consistent across two important cross-sections of the 30-trait morphospace: the two discriminant axes separating the three parental species (Fig. 3, Table S2) and the two major axes of nonlinear selection estimated from generalized projection pursuit regression (Fig. 4, Tables S3-S4). The lack of any signal of frequency-dependence suggests that some features of the survival fitness landscape are robust to competitive environment and reflect intrinsic viability or organismal performance constraints.

Combined estimates of the survival fitness landscape indicated a surprisingly consistent topography across both independent field experiments (Fig. 5: 2014/2015 and 2011) comprised of an isolated fitness peak corresponding to the generalist phenotype separated by a small fitness valley from a higher fitness peak corresponding to the molluscivore phenotype. For survival, multiple peaks emerged from evidence of higher survival of generalist and molluscivore phenotypes after three months in lake 1 combined with evidence of much higher survival of the molluscivore phenotype after 11 months in lake 2 (Fig. 3). Growth rate in lake 1 also showed evidence of multiple peaks due to higher growth rates of generalist phenotypes in both enclosures combined with moderate molluscivore growth rates and very low scale-eater growth rates in the low-frequency enclosure (Fig. S7). These joint landscapes inferred from the best-fitting models are admittedly reflective of combining different frequency treatments, exposure periods, lake environments, and potentially different selective regimes; however, formal comparisons of general additive models provided no evidence of different selective regimes between treatments and only a fixed effect of lake environment, rather than a change in fitness landscape topography between lake environments (e.g. Table S6: very low support for models including a ‘by lake’ effect), supporting inference of a single combined selective environment. Furthermore, our interest is in the combined landscape driving the adaptive radiation of species differences across lake environments, rather lake-specific local adaptive regimes unrelated to species divergence.

Across all four field enclosures and both fitness proxies the most consistent feature of fitness landscape topography was a large fitness valley isolating hybrids resembling the scale-eater, the most morphologically, ecologically, and genetically divergent specialist in the radiation, from all other hybrids across treatments, lake environments, field exposure periods, independent field experiments, and across nearly all dimensions of the 30-trait hybrid morphospace (Figs. 3-6). This persistent multi-peak fitness landscape within San Salvador Island’s hypersaline lakes provides a striking explanation for the rarity of trophic specialization across the Caribbean if outgroup generalist populations are isolated on local fitness optimum and experiencing stabilizing selection opposing trait diversification (Martin & Wainwright 2013a).

**Fig. 6.**
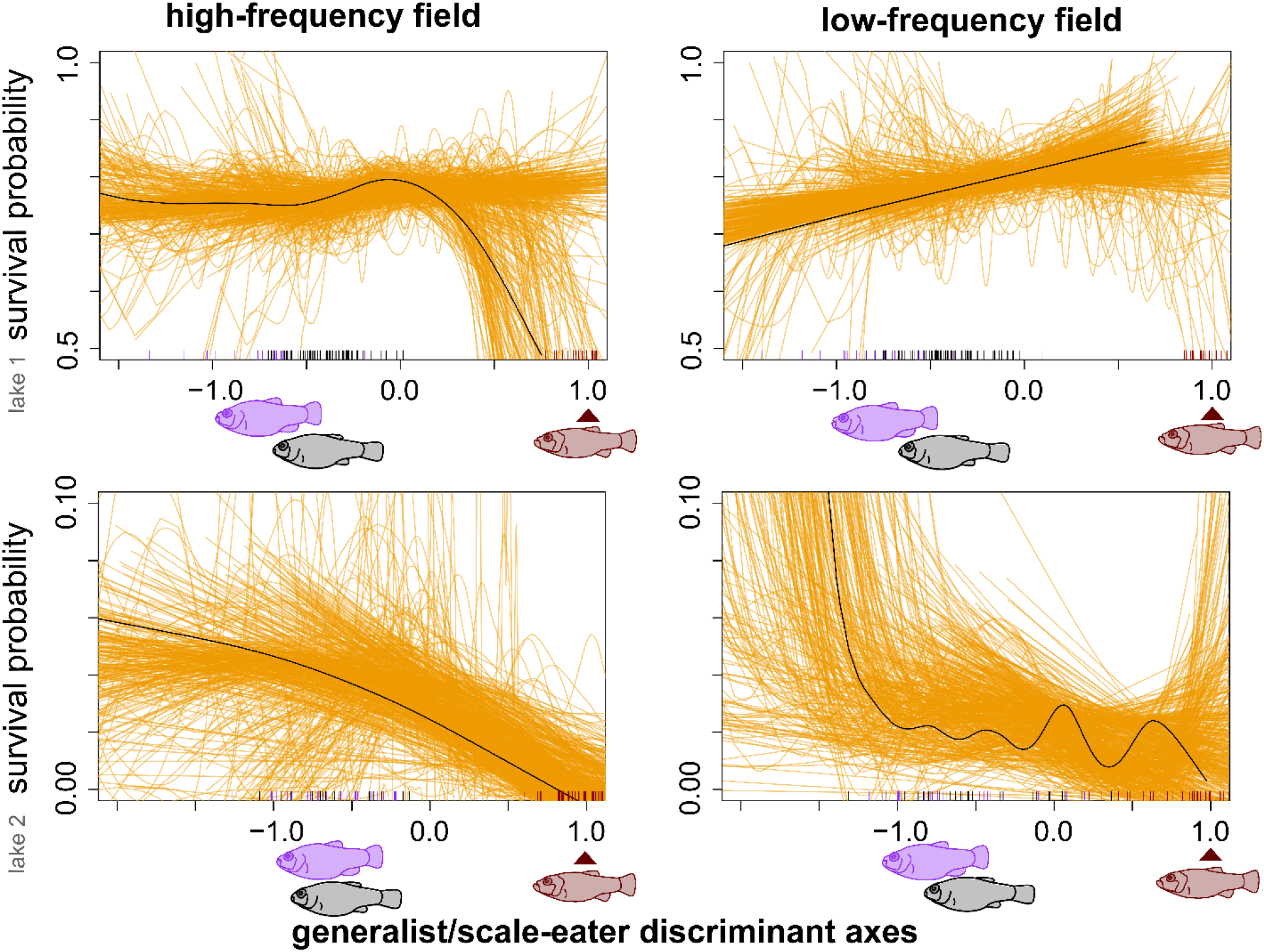
Spaghetti plots illustrate all possible fitness paths between generalist and scale-eater hybrid phenotypes in high- (first column) and low-frequency (second column) field enclosures. Each orange line depicts the relationship between survival and a random discriminant axis separating generalist and scale-eater phenotypes estimated from generalized cross-validation of a smoothing spline for 500 random subsets of 15 size-corrected traits (out of 30); the black line illustrates the smoothing spline estimated for a discriminant axis from all 30 traits, the grey lines illustrate the smoothing spline estimated for each subset. Each subsampled discriminant axis was rescaled so that the mean parental scale-eater phenotype = 1 (red arrows). Parental phenotypes are illustrated as black (generalist), purple (molluscivore), and red (scale-eater) tick marks on the x-axis.

### Extrinsic and intrinsic factors could explain low survival of transgressive hybrid phenotypes

In contrast to predictions of negative frequency-dependent disruptive selection (Gigord *et al.* 2001; Bolnick 2004; Haller & Hendry 2014), rare and transgressive hybrid phenotypes outside parental ranges exhibited the lowest survival rates within field enclosures. One possible explanation is that the intrinsic viability and performance of survival tasks by these hybrids was impaired due to their mosaic hybrid genetic backgrounds. This could result in mismatched craniofacial and behavioral traits leading to poor foraging performance. For example, the large oral jaws of scale-eaters appear to result from at least four moderate effect quantitative trait loci on different linkage groups that each increase jaw size (Martin *et al.* 2017), suggesting that the genetic basis of even this single trait is moderately polygenic and may not be fully recovered within F4/F5 hybrids. Furthermore, F1 hybrid scale-eaters exhibit foraging kinematics during scale-biting strikes similar to generalists, suggesting that kinematic behaviors may be non-additive and severely mismatched in more advanced hybrids (St. John & Martin 2019). Impaired foraging performance of transgressive hybrids in the field environment is further supported by the observation that hybrids with scale-eater morphologies in laboratory control aquaria exhibited the highest survival rates when fed only pellet foods (Figs. 3-4), demonstrating that the field environment is contributing to the low survival of scale-eater hybrids. This hypothesis is consistent with laboratory studies of feeding performance in other hybrid fishes (Parnell *et al.* 2008; Arnegard *et al.* 2014; Matthews & Albertson 2017; Selz & Seehausen 2019).

An alternative and non-mutually exclusive possibility is that intrinsic genetic incompatibilities in hybrids are contributing to their low survival rates, particularly if more transgressive hybrid phenotypes are associated with a greater number or more severe genetic incompatibilities. Although the San Salvador radiation only diverged approximately 10,000 years old, trophic specialists within the radiation contain ancient adaptive variants also found in outgroups that diverged over 5 million years ago (Richards et al. in prep.). Genetic incompatibility loci are known to segregate in wild populations ((Fishman *et al.* 2013); reviewed in (Cutter 2012)) and hundreds of genetic incompatibility loci have been found in hybrid zones between swordtail fish species (Schumer *et al.* 2014; Schumer & Brandvain 2016). In support of this hypothesis, F1 hybrids of specialist species within the San Salvador radiation show evidence of hybrid gene misregulation in approximately 10% of their differentially expressed genes, i.e. transgressive gene expression levels significantly different from parental expression levels, in whole larvae at 8 days post fertilization (dpf) and within craniofacial tissues at 17-20 dpf, respectively (McGirr & Martin 2019b, a). Although the fitness effects on hybrids are unknown, hybrid gene misregulation has been shown to affect hybrid viability and sterility in other systems (Ortiz-Barrientos *et al.* 2007; Renaut *et al.* 2009; Renaut & Bernatchez 2011; Mack *et al.* 2016; Mack & Nachman 2017) and misregulated genes in San Salvador Island pupfish species are enriched for developmental processes affecting ecological traits relevant to trophic specialization, including craniofacial morphology, muscle mass, and nitrogen metabolism (McGirr & Martin 2019a). However, the link between transgressive hybrid phenotypes, divergent ecological selection, and the extent of gene misregulation or other genetic incompatibilities is still unknown (Kulmuni & Westram 2017).

### Biomechanical constraints of scale-eating suggest a large and stable fitness valley isolates this novel ecological niche

The rare evolution, high performance demands, and low caloric payoffs of scale-eating suggest that a wide and deep fitness valley isolates this niche and the necessary adaptive traits from all other ecological niches. Scale-eating (lepidophagy) is a particularly rare trophic niche among fishes and has evolved independently only 19 times across diverse marine, coastal, riverine, and lacustrine environments (Sazima 1983; Martin & Wainwright 2013b; Kolmann *et al.* 2018; St. John *et al.* 2018) and ontogenetic stages from only juveniles to obligate scale-eating adults (Grubh & Winemiller 2004; Janovetz 2005; Koblmüller *et al.* 2007; Raffini & Meyer 2018). In particular, it only evolved once in Cyprinodontiformes within the San Salvador Island radiation and is thus separated by 168 million years of evolutionary time from the most closely related African cichlid scale-eating specialists (Martin & Wainwright 2013b). Second, scale-eating pupfish strike frequently in the wild, approximately once per minute, resulting in only a few scales and mouthful of skin mucus per high-speed strike completed within 10 - 15 ms (St. John & Martin 2019a). Thus, the energetic payoff from scales is most likely low relative the energetic demands of high-speed scale-eating or scale-rasping strikes, consistent with the observation that all scale-eaters are size-limited relative to their prey, unlike piscivorous fishes which generally grow much larger (Sazima *et al.* 1983).

Consistent with these biomechanical predictions of a large fitness valley, our empirical field fitness data indicate that hybrids resembling scale-eaters suffered the highest fitness costs in growth and survival across all treatments, lakes, trait subsets, time periods, and across independent field experiments but not in laboratory control aquaria while fed only pellet foods. This provides an unexpected explanation for the rarity of scale-eating across Cyprinodontiform fishes if adapting to this specialized trophic niches requires multiple phenotypic traits that only provide fitness benefits in combination (i.e. fitness epistasis (Whitlock *et al.* 1995)). This is also supported by existing population genomic and quantitative genetic evidence which suggests that adaptation to scale-eating is multifactorial and that multiple sources of adaptive variation from across the Caribbean contributed to trait diversification in this radiation (McGirr & Martin 2016; Martin *et al.* 2017; Richards & Martin 2017). For example, aggressive and energetically demanding scale-eating behaviors may be highly deleterious in individuals with small oral jaws (St. John & Martin 2019b). Conversely, algae-scraping with enlarged oral jaws or highly energetic strikes by hybrids resembling the generalist may also not provide a sufficient energy surplus for survival under field conditions.

## Conclusion: on the origins of novelty during adaptive radiation on multiple fitness peaks

The sensitivity of fitness landscape topography to the environment is rarely measured beyond a single population or fitness peak. Here we experimentally tested the effect of competitor frequency on a fitness surface spanning diverse hybrid phenotypes within a nascent adaptive radiation, comparable to phenotypic divergence spanning over 20 million years of Cyprinodontiform evolution. Hybrid survival showed no signal of frequency-dependence, challenging existing competitive speciation theory and many previous experiments conducted within a single population or species pair. Furthermore, major features of these complex fitness landscapes were shared across different lake environments, competitor frequencies, and independent field experiments: multiple fitness peaks and a large fitness valley isolating the scale-eating phenotype. This challenges our existing view of empirical fitness surfaces as highly sensitive to environmental perturbation. Instead, multi-peak fitness landscapes spanning macroevolutionary levels of phenotypic disparity display both static and dynamic features across space, time, and competitive environment. These empirical results strengthen the connection between microevolutionary dynamic processes and macroevolutionary patterns of stasis by investigating the complex interplay of organismal phenotype and environment within a rapidly diversifying radiation.

## Acknowledgments

We thank the Miller Institute for Basic Research in Science, the University of California, Berkeley, the University of North Carolina at Chapel Hill, NSF CAREER award 1749764, and NIH 5R01DE027052-02 for funding to CHM. The Bahamas Environmental Science and Technology Commission and the Ministry of Agriculture kindly provided permission to export, import, and tag fish and conduct this research. Erica Bree Rosenblum generously provided laboratory animal space at the University of California, Berkeley. Rochelle Hanna, Velda Knowles, and the Gerace Research Centre provided logistical assistance; Clare Bocklage, Alexander Payne, Christina Lim, Qiongqiong Mei, Ivan Piedad, Sarah Bencuya, Stephanie Jeselson, Courtney Farge, Kristi Dixon, and David Richard assisted with morphological measurements; and Erica Bree Rosenblum, Craig Miller, Emilie Richards, Michelle St. John, and Joseph McGirr provided helpful comments on the project. All animal care protocols were approved by the University of California, Berkeley and the University of North Carolina at Chapel Hill Animal Care and Use Committees.

## Data Accessibility Statement

All data used for this study will be deposited in the Dryad Digital Repository upon acceptance of the manuscript.

## Ethics Statement

The Bahamas Environmental Science and Technology Commission and the Ministry of Agriculture kindly provided permission to export, import, and tag fish and conduct this research (research permits renewed annually from 2011 – 2015 at the conclusion of this study). All animal care protocols were approved by the University of California, Berkeley and the University of North Carolina at Chapel Hill Animal Care and Use Committees.

## Author contributions

CHM designed the study, performed field experiments, analyzed the data, wrote and edited the manuscript, and funded the study. KG contributed substantially to laboratory data collection.

## References

Abrams, P. (2001). Modelling the adaptive dynamics of traits involved in inter- and intarspecific interactions: An assesment of three methods. Ecol. Lett., 4, 166–175.

Abrams, P., Matsuda, H. & Harada, Y. (1993). Evolutionarily unstable fitness maxima and stable fitness minima of continuous traits. Evol. Ecol.

Arnegard, M.E., McGee, M.D., Matthews, B., Marchinko, K.B., Conte, G.L., Kabir, S., et al. (2014). Genetics of ecological divergence during speciation. Nature, 511, 307–311.

Arnold, S.J., Pfrender, M.E. & Jones, A.G. (2001). The adaptive landscape as a conceptual bridge between micro- and macroevolution. Genetica, 112–113, 9–32.

Baptestini, E.M., de Aguiar, M. a M., Bolnick, D.I. & Araújo, M.S. (2009). The shape of the competition and carrying capacity kernels affects the likelihood of disruptive selection. J. Theor. Biol., 259, 5–11.

Beaulieu, J. & O’Meara, B. (2012). OUwie: an analysis of evolutionary rates in an OU framework.

Bolnick, D.I. (2004). Can Intraspecific Competition Drive Disruptive Selection◻? an Experimental Test in Natural Populations of Sticklebacks Can Intraspecific Competition Drive Disruptive Selection◻? an Experimental Test in Natural Populations of Sticklebacks. Evolution (N. Y)., 58, 608–618.

Bolnick, D.I. & Lau, O.L. (2008). Predictable patterns of disruptive selection in stickleback in postglacial lakes. Am. Nat., 172, 1–11.

Bolnick, D.I. & Stutz, W.E. (2017). Frequency dependence limits divergent evolution by favouring rare immigrants over residents. Nature, 546, 285–288.

Boucher, F. & Demery, V. (2016). Inferring bounded evolution in phenotypic characters from phylogenetic comparative data. Syst. Biol., In press.

Bürger, R. & Schneider, K.A. (2006). Intraspecific competitive divergence and convergence under assortative mating. Am. Nat., 167, 190–205.

Butler, M.A. & King, A.A. (2004). Phylogenetic comparative analysis: a modeling approach for adaptive evolution. Am. Nat., 164, 683–695.

Calsbeek, R., Bonvini, L. & Box, R. (2009). GEOGRAPHIC VARIATION, FREQUENCY-DEPENDENT SELECTION, AND THE MAINTENANCE OF A FEMALE-LIMITED POLYMORPHISM. Evolution (N. Y)., 64, 116–125.

Carneiro, M. & Hartl, D.L. (2010). Adaptive landscapes and protein evolution. Proc. Natl. Acad. Sci., 107, 1747–1751.

Cutter, A.D. (2012). The polymorphic prelude to Bateson-Dobzhansky-Muller incompatibilities. Trends Ecol. Evol., 27, 209–18.

Dieckmann, U. & Doebeli, M. (1999). On the origin of species by sympatric speciation. Nature, 400, 354–7.

Doebeli, M. & Dieckmann, U. (2005). Adaptive dynamics as a mathematical tool for studying the ecology of speciation processes. J. Evol. Biol., 18, 1194–200.

Doebeli, M., Dieckmann, U., Metz, J.A. & Tautz, D. (2005). What we have also learned: adaptive speciation is theoretically plausible. Evolution (N. Y)., 59, 691–699.

Enquist, B. & Niklas, K. (2002). Global allocation rules for patterns of biomass partitioning in seed plants. Science (80-.)., 22, 1571–1520.

Fisher, R. (1930). Genetical theory of natural selection. Clarendon Press.

Fishman, L., Stathos, A., Beardsley, P.M., Williams, C.F. & Hill, J.P. (2013). Chromosomal rearrangements and the genetics of reproductive barriers in mimulus (monkey flowers). Evolution (N. Y)., 67, 2547–2560.

Furness, A. & Reznick, D. (2015). Convergent evolution of alternative developmental trajectories associated with diapause in African and South American killifish. Proceeding R. Soc. London Ser. B.

Gavrilets, S. (1997). Evolution and speciation on holey adaptive landscapes. Trends Ecol. Evol., 12, 307–312.

Gavrilets, S. (1999). A Dynamical Theory of Speciation on Holey, 154.

Gavrilets, S. (2004). Fitness landscapes and the origin of species. Princeton University Press.

Gigord, L.D., Macnair, M.R. & Smithson, a. (2001). Negative frequency-dependent selection maintains a dramatic flower color polymorphism in the rewardless orchid Dactylorhiza sambucina (L.) Soo. Proc. Natl. Acad. Sci. U. S. A., 98, 6253–5.

Grant, P.R. & Grant, B.R. (2002). Unpredictable Evolution in a 30-Year Study of Darwin ’ s Finches, 296, 707–712.

Grubh, A.R. & Winemiller, K.O. (2004). Ontogeny of Scale Feeding in the Asian Glassfish, Chanda nama (Ambassidae). Copeia, 2004, 903–907.

Haller, B.C. & Hendry, A.P. (2014). Solving the paradox of stasis: squashed stabilizing selection and the limits of detection. Evolution (N. Y)., 68, 483–500.

Hansen, T.F., Pienaar, J. & Orzack, S.H. (2008). A comparative method for studying adaptation to a randomly evolving environment. Evolution, 62, 1965–77.

Harmon, L., Andreazzi, C., Debarre, F., Drury, J., Goldberg, E. & Martins, A. (2019). Detecting the Macroevolutionary Signal of Species Interactions. J. Evol. Biol., In press.

Harmon, L.J., Losos, J.B., Davies, T.J., Gillespie, R.G., Gittleman, J.L., Jennings, W.B., et al. (2010). Early bursts of body size and shape evolution are rare in comparative data. Evolution, 64, 2385–2396.

Higham, T.E., Rogers, S.M., Langerhans, R.B., Jamniczky, H.A., Lauder, G. V., Stewart, W.J., et al. (2016). Speciation through the lens of biomechanics: locomotion, prey capture and reproductive isolation. Proc. R. Soc. B Biol. Sci., 283, 20161294.

Hori, M. (1993). Frequency-Dependent Natural-Selection in the Handedness of Scale-Eating Cichlid Fish. Science (80-.)., 260, 216–219.

Janovetz, J. (2005). Functional morphology of feeding in the scale-eating specialist Catoprion mento. J. Exp. Biol., 208, 4757–4768.

St. John, M. & Martin, C. (2019a). Scale-eating specialists evolved adaptive feeding kinematics within a microendemic radiation of San Salvador Island pupfishes. bioRxiv.

St. John, M.E. & Martin, C.H. (2019b). Scale-eating specialists evolved adaptive feeding kinematics within a microendemic radiation of San Salvador Island pupfishes. bioRxiv, 648451.

St. John, M.E., McGirr, J.A. & Martin, C.H. (2018). The behavioral origins of novelty: did increased aggression lead to scale-eating in pupfishes? Behav. Ecol.

Kassen, R., Llewellyn, M. & Rainey, P.B. (2004). Ecological constraints on diversification in a model adaptive radiation. Nature, 431, 984–988.

Keagy, J., Lettieri, L. & Boughman, J.W. (2015). Male competition fitness landscapes predict both forward and reverse speciation. Ecol. Lett., 19, 71–80.

Kingsolver, J.G., Hoekstra, H.E., Hoekstra, J.M., Berrigan, D., Vignieri, S.N., Hill, C.E., et al. (2001). The strength of phenotypic selection in natural populations. Am. Nat., 157, 245–61.

Koblmüller, S., Egger, B., Sturmbauer, C. & Sefc, K.M. (2007). Evolutionary history of Lake Tanganyika’s scale-eating cichlid fishes. Mol. Phylogenet. Evol., 44, 1295–305.

Kolmann, M.A., Huie, J.M., Evans, K. & Summers, A.P. (2018). Specialized specialists and the narrow niche fallacy: A tale of scale-feeding fishes. R. Soc. Open Sci., 5.

Kopp, M., Servedio, M.R., Mendelson, T.C., Safran, R.J., Rodríguez, R.L., Hauber, M.E., et al. (2017). Mechanisms of Assortative Mating in Speciation with Gene Flow: Connecting Theory and Empirical Research. Am. Nat., 191, 000–000.

Koskella, B. & Lively, C.M. (2009). Evidence for negative frequency-dependent selection during experimental coevolution of a freshwater snail and a sterilizing trematode. Evolution, 63, 2213–21.

Kulmuni, J. & Westram, A.M. (2017). Intrinsic incompatibilities evolving as a by-product of divergent ecological selection: Considering them in empirical studies on divergence with gene flow. Mol. Ecol., 26, 3093–3103.

Kusche, H., Lee, H.J. & Meyer, A. (2012). Mouth asymmetry in the textbook example of scale-eating cichlid fish is not a discrete dimorphism after all Subject collections Mouth asymmetry in the textbook example of scale-eating cichlid fish is not a discrete dimorphism after all.

Lande, R. (1979). Quantitative genetic analysis of multivariate evolution, applied to brain: body size allometry. Evolution, 33, 402–416.

Lande, R. & Arnold, S.J. (1983). The Measurement of Selection on Correlated Characters. Evolution (N. Y)., 37, 1210.

Landis, M., Edwards, E. & Donoghue, M. (2019). Modeling phylogenetic biome shifts on a planet with a past. bioRxiv, https://doi.org/10.1101/832527.

Lavin, A. & McPhail, J. (1986). Adaptive Divergence of Trophic Phenotype among Freshwater Populations of the Threespine Stickleback (Gasterosteus aculeatus). Can. J. Fish. Aquat. Sci., 43, 2455–2463.

Mack, K., Campbell, P. & Nachman, M. (2016). Gene regulation and speciation in house mice. Genome Res., 26, 451–461.

Mack, K. & Nachman, M. (2017). Gene regulation and speciation. Trends Genet., 33, 68–80.

Martin, C.H. (2016a). Context dependence in complex adaptive landscapes◻: frequency and trait-dependent selection surfaces within an adaptive radiation of Caribbean pupfishes. Evolution (N. Y)., 1–18.

Martin, C.H. (2016b). The cryptic origins of evolutionary novelty: 1000◻fold faster trophic diversification rates without increased ecological opportunity or hybrid swarm. Evolution (N. Y)., 70.11, 2504–2519.

Martin, C.H., Crawford, J.E., Turner, B.J. & Simons, L.H. (2016). Diabolical survival in Death Valley: recent pupfish colonization, gene flow, and genetic assimilation in the smallest species range on earth. Proc. R. Soc. B Biol. Sci., 283, 23–34.

Martin, C.H., Erickson, P.A. & Miller, C.T. (2017). The genetic architecture of novel trophic specialists: larger effect sizes are associated with exceptional oral jaw diversification in a pupfish adaptive radiation. Mol. Ecol., 26, 624–638.

Martin, C.H. & Feinstein, L.C. (2014). Novel trophic niches drive variable progress towards ecological speciation within an adaptive radiation of pupfishes. Mol. Ecol., 23, 1846–62.

Martin, C.H. & Richards, E.J. (2019). The paradox behind the pattern of rapid adaptive radiation: how can the speciation process sustain itself through an early burst? Annu. Rev. Ecol. Evol. Syst., In press.

Martin, C.H. & Wainwright, P.C. (2011). Trophic novelty is linked to exceptional rates of morphological diversification in two adaptive radiations of Cyprinodon pupfishes. Evolution, 65, 2197–212.

Martin, C.H. & Wainwright, P.C. (2013a). Multiple fitness peaks on the adaptive landscape drive adaptive radiation in the wild. Science, 339, 208–211.

Martin, C.H. & Wainwright, P.C. (2013b). On the measurement of ecological novelty: scale-eating pupfish are separated by 168 my from other scale-eating fishes. PLoS One, 8, e71164.

Matessi, C., Gimelfarb, A. & Gavrilets, S. (2002). Long-term buildup of reproductive isolation promoted by disruptive selection: how far does it go? Selection, 2, 41–64.

Matthews, D.G. & Albertson, R.C. (2017). Effect of craniofacial genotype on the relationship between morphology and feeding performance in cichlid fishes. Evolution (N. Y)., 71, 2050–2061.

McGirr, J. & Martin, C. (2019a). Ecological divergence in sympatry causes gene misregulation in hybrids. bioRxiv, 717025.

McGirr, J. & Martin, C. (2019b). Hybrid misexpression in multiple developing tissues within a recent adaptive radiation of Cyprinodon pupfishes. bioRxiv, 372912.

McGirr, J.A. & Martin, C.H. (2016). Novel candidate genes underlying extreme trophic specialization in Caribbean pupfishes. Mol. Biol. Evol., msw286.

Mitchell-Olds, T. & Shaw. (1987). Regression analysis of natural selection: statistical inference and biological interpretation. Evolution (N. Y)., 41, 1149–1161.

Morrissey, M.B. (2014). In search of the best methods for multivariate selection analysis. Methods Ecol. Evol., n/a–n/a.

Nosil, P., Villoutreix, R., de Carvalho, C.F., Farkas, T.E., Soria-Carrasco, V., Feder, J.L., et al. (2018). Natural selection and the predictability of evolution inTimemastick insects. Science (80-.)., 359, 765–770.

Nychka, D., Furrer, R., Paige, J. & Sain, S. (2017). fields: Tools for spatial data. R Packag. version 9.6.

O’Meara, B.C. (2012). Evolutionary Inferences from Phylogenies: A Review of Methods. Annu. Rev. Ecol. Evol. Syst., 43, 267–285.

Olendorf, R., Rodd, F.H., Punzalan, D., Houde, A.E., Hurt, C., Reznick, D.N., et al. (2006). Frequency-dependent survival in natural guppy populations. Nature, 441, 633–6.

Ortiz-Barrientos, D., Counterman, B. & Noor, M. (2007). Gene expression divergence and the origin of hybrid dysfunctions. Genetica, 129, 71–81.

Otto, S.P., Servedio, M.R. & Nuismer, S.L. (2008). Frequency-dependent selection and the evolution of assortative mating. Genetics, 179, 2091–112.

Parnell, N.F., Hulsey, C.D. & Streelman, J.T. (2008). Hybridization produces novelty when the mapping of form to function is many to one. BMC Evol. Biol., 8, 122.

Pfennig, D. (1992). POLYPHENISM IN SPADEFOOT TOAD TADPOLES AS A LOCALLY ADJUSTED EVOLUTIONARILY STABLE STRATEGY. Evolution (N. Y)., 46, 1408–1420.

Polechová, J. & Barton, N.H. (2005). Speciation Through Competition◻: a Critical Review Speciation Through Competition◻: a Critical Review, 59, 1194–1210.

Rabosky, D.L. (2009). Ecological Limits on Clade Diversification in Higher Taxa. Am. Nat., 173, 662–674.

Raffini, F. & Meyer, A. (2018). A comprehensive overview of the developmental basis and adaptive significance of a textbook polymorphism: head asymmetry in the cichlid fish Perissodus microlepis. Hydrobiologia, 0123456789.

Renaut, S. & Bernatchez, L. (2011). Transcriptome-wide signature of hybrid breakdown associated with intrinsic reproductive isolation in lake whitefish species pairs (Coregonus spp. Salmonidae). Heredity (Edinb)., 106, 1003–1011.

Renaut, S., Nolte, A. & Bernatchez, L. (2009). Gene expression divergence and hybrid misexpression between lake whitefish species pairs (Coregonus spp. Salmonidae). Mol. Biol. Evol., 26, 925–936.

Richards, E.J. & Martin, C.H. (2017). Adaptive introgression from distant Caribbean islands contributed to the diversification of a microendemic adaptive radiation of trophic specialist pupfishes. PLoS Genet., 13, 1–35.

Sazima, I. (1983). Scale-eating in characoids and other fishes. Environ. Biol. Fishes, 9, 87–101.

Sazima, I., Zoologia, D. De, Campinas, U.E. De & Paulo, S. (1983). Scale-eating in characoids and other fishes, 9.

Schluter, D. (2003). Frequency dependent natural selection during character displacement in sticklebacks. Evolution, 57, 1142–50.

Schluter, D., Columbia, B. & E-mail, C. (2003). Frequency dependent natural selection during character displacement in sticklebacks. Evolution, 57, 1142–50.

Schluter, D. & Nychka, D. (1994). Exploring fitness surfaces. Am. Nat., 143, 597–616.

Schumer, M. & Brandvain, Y. (2016). Determining epistatic selection in admixed populations. Mol. Ecol., 25, 2577–2591.

Schumer, M., Cui, R., Powell, D.L., Dresner, R., Rosenthal, G.G. & Andolfatto, P. (2014). High-resolution mapping reveals hundreds of genetic incompatibilities in hybridizing fish species. Elife, 3, 1–21.

Seehausen, O. & Schluter, D. (2004). Male-male competition and nuptial-colour displacement as a diversifying force in Lake Victoria cichlid fishes. Proc. Biol. Sci., 271, 1345–53.

Selz, O. & Seehausen, O. (2019). Interspecific hybridization can generate functional novelty in cichlid fish. Proceeding R. Soc. London Ser. B, 286.

Servedio, M.R. & Burger, R. (2014). The counterintuitive role of sexual selection in species maintenance and speciation. Proc. Natl. Acad. Sci., 111, 8113–8118.

Simpson, G. (1944a). Tempo and mode in evolution.

Simpson, G.G. (1944b). Tempo and Mode in Evolution, 197–217.

Sinervo, B., Svensson, E. & Comendant, T. (2000). Density cycles and an offspring quantity and quality game driven by natural selection. Nature, 406, 985–8.

Svensson, E.I. & Calsbeek, R. (2012). The adaptive landscape. Oxford University Press, Oxford.

Uyeda, J.C. & Harmon, L.J. (2014). A novel Bayesian method for inferring and interpreting the dynamics of adaptive landscapes from phylogenetic comparative data. Syst. Biol., 63, 902–918.

Weeks, A. & Hoffmann, A. (2008). Frequency-dependent selection maintains clonal diversity in an asexual organism. Proc. Natl. Acad. Sci. USA, 105, 17872–17877.

Weinreich, D.M., Watson, R. a & Chao, L. (2005). Perspective: Sign epistasis and genetic constraint on evolutionary trajectories. Evolution, 59, 1165–74.

Whitlock, M., Phillips, P., Moore, F. & Tonsor, S. (1995). Multiple fitness peaks and epistasis. Annu. Rev. Ecol. Syst., 26, 601–629.

Wright, S. (1932). The roles of mutation, inbreeding, crossbreeding and selection in evolution. Proc. Sixth Int. Congr. Genet., 1, 356–366.

